# Ghrelin and relamorelin alleviate hypoglycaemia in humanised mice with congenital hyperinsulinism

**DOI:** 10.1101/2025.10.02.680116

**Authors:** Väinö Lithovius, Hossam Montaser, Jonna Saarimäki-Vire, Hazem Ibrahim, Tom Barsby, Diego Balboa, Timo Otonkoski

## Abstract

K_ATP_-channel related hyperinsulinism (K_ATP_HI) is a genetic disorder of the pancreatic beta cells, manifesting as life-threatening hypoglycaemia in neonates due to dysregulated insulin secretion. Management of the severe diffuse form of K_ATP_HI currently lacks treatment options, as the first-line therapy octreotide is often insufficiently effective, necessitating radical pancreatectomy in many patients. In this study, we applied stem cell derived islets carrying K_ATP_HI-causing mutation *KCNJ11*^-/-^ to develop new pharmaceutical therapies for K_ATP_HI. We tested seven candidate molecules *in vitro*, identifying three that reduced insulin secretion in *KCNJ11*^-/-^ stem cell derived islets. Of these, we tested acyl-ghrelin and its long-acting analogue relamorelin in mice that became hypoglycaemic due to carrying human *KCNJ11*^-/-^ stem cell derived grafts. Acyl-ghrelin and relamorelin alone increased fasting glycaemia. Combining relamorelin with octreotide increased blood glucose more than the sum of each drug alone, reversing hypoglycaemia to normoglycaemia. In conclusion, we show that ghrelin receptor agonists have acute anti-hypoglycaemic effects in a humanised mouse model of K_ATP_HI, especially when combined with octreotide. Relamorelin has been tested in >650 diabetic adults with little side effects, leading us to propose relamorelin – octreotide combination as a novel therapeutic candidate for severe K_ATP_HI patients.

## INTRODUCTION

Maintenance of correct glucose level, euglycaemia, is critical for human health. Prevention of hypoglycaemia is especially important, as plasma glucose < 1 mmol/l is acutely life-threatening^1^. Congenital hyperinsulinism (CHI) encompasses inborn defects of pancreatic beta cells that cause inappropriate insulin hypersecretion, which manifests as hypoglycaemia usually in the neonatal period. While at least 15 genes have been identified as causative for CHI, more than 50 % of cases are caused by loss-of-function mutations in genes encoding the beta cell K_ATP_-channel, *KCNJ11* and *ABCC8*, termed K_ATP_HI ^2,3^. K_ATP_HI is a distinct clinical entity due to its severity and its two histological subtypes, the focal and diffuse forms ^4^. Positron emission tomography guided surgery presents a curative treatment for the focal form ^5,6^. However, management of the diffuse form, where all beta cells are affected remains challenging.

The molecular mechanism of diffuse K_ATP_HI makes it especially challenging to manage. Closure of the K_ATP_-channel is the key trigger of insulin secretion ^7^ and its dysfunction exerts a profound influence on beta cell function. K_ATP_HI causing mutations also render the channel inaccessible to pharmacotherapy with diazoxide, whose active site is on the missing K_ATP_-channels ^8,9^. The therapeutic mainstay is thus somatostatin receptor (SSTR) agonists, such as octreotide ^10,11^, complemented by agents such as glucagon, calcium channel blockers and diuretics and supportive treatments like glucose infusions ^4,12^.

The current state-of-the-art management of K_ATP_HI is suboptimal: the aforementioned therapy is effective in some patients in stabilising severe hypoglycaemia, but leaves a risk for learning difficulties in the long term ^13,14^. For many patients, surgical intervention in the form of near-total pancreatectomy is the only option. This radical procedure is associated with lifelong diabetes and exocrine insufficiency, without completely eliminating the risk of continued hypoglycaemia ^5,15^. While usually effective as an anti-hypoglycaemic agent, octreotide therapy is associated with around 1-2% chance for necrotising enterocolitis, a potentially fatal side effect ^16–18^. Thus, the development of new pharmaceuticals, such as long-acting glucagon analogues or GLP1-receptor antagonists is ongoing ^19–21^. Nevertheless, there is still a need to find more effective agents for improved pharmacological treatment of K_ATP_HI, to reduce the need for near-total pancreatectomy and prevent brain damage in the short and long term.

This pursuit is hampered by limitations in modelling systems, as K_ATP_HI patient primary islets are scarce and transgenic rodent models do not recapitulate the severity of human K_ATP_HI ^22,23^. In this study, we used *KCNJ11*^-/-^ stem-cell-derived islets (SC-islets), which present important advantages over the other modelling systems. SC-islets can be produced in high quantities with high consistency ^24–26^ and recapitulate human human beta cell insulin secretion function accurately ^27^. We present here a humanised mouse model of K_ATP_HI based on transplanted SC-islets and identify ghrelin receptor agonists as a potential new class of anti-hypoglycaemic drugs for improving pharmaceutical therapy of K_ATP_HI.

## RESULTS

### KCNJ11^-/-^ stem cell derived islets have K_ATP_HI phenotype in vitro

We generated an embryonic stem cell line with a knockout of the *KCNJ11* gene ^28^). *KCNJ11* encodes the pore-forming subunit of the K_ATP_-channel, whose loss results in K_ATP_HI due the absence of hyperpolarization typically established in beta cells by open K_ATP_-channels under low glucose conditions. Into the same embryonic stem cell line (H1), we generated a *KCNJ11*^R201H^ mutation ^28^, causative for K_ATP_-channel -related neonatal diabetes ^29^. We differentiated these *KCNJ11*-mutant stem cells along with unedited *KCNJ11*^+/+^ stem cells to stem cell derived islets (SC-islets) following a state-of-the art protocol with seven stages that produces SC-islets with glucose responsive beta cells by mimicking human islet development *in vitro* ^27,30^.

To assess how closely the SC-islets recapitulate the K_ATP_HI and diabetic phenotypes, we studied their insulin secretion weekly during the last stage (stage 7, S7) of the differentiation, a stage we have previously shown to induce maturation of insulin secretion function ^27^. Insulin secretion at low glucose levels (3 mM, sub-stimulatory for healthy beta cells) was higher in *KCNJ11*^-/-^ SC-islets compared to *KCNJ11*^+/+^ SC-islets from S7w2 onwards. This difference progressively increased, with *KCNJ11*^-/-^ cells exhibiting approximately fourfold higher basal insulin secretion by S7w6 (Fig. 1a). This represents a greater difference between the healthy and K_ATP_HI cells compared to our previous SC-islet model, which was based on *ABCC8*-mutant SC-islets differentiated with previous generation SC-islet differentiation protocols ^31^.

**Figure 1.**
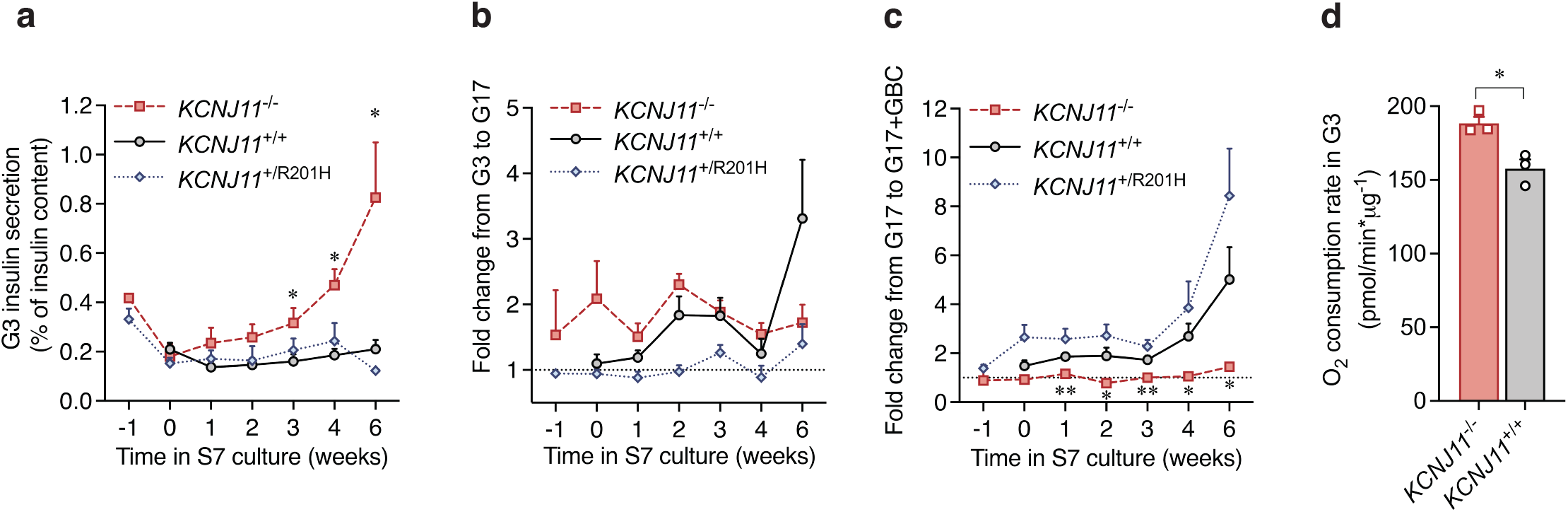
*in vitro* phenotype of the *KCNJ11*-variant SC-islets: **a-c)** Data from sequential static incubations in 30 min each of 3 mmol/l glucose (G3), 17 mmol/l glucose (G17) and G17 + 100 nmol/l glibenclamide (GBC), tested weekly in stage 7 (S7) culture and at the start of stage 6 culture (-1 timepoint). Absolute G3 insulin release in a), relative change from G3 to G17 in b) and from G17 to G17+GBC in c). Significance against *KCNJ11*^+/+^ tested with multiple Welch’s t-tests. **d)** Respiration rate in G3, normalised to DNA, tested at S7w6. Welch’s t-test.

When stimulated with glucose, the *KCNJ11*^-/-^ SC-islets maintained a low response, while the *KCNJ11*^+/+^ controls became strongly glucose responsive by S7w6 ^27^. As expected, the *KCNJ11*^R201H^ SC-islets remained unresponsive to glucose (Fig. 1b). The *KCNJ11*^-/-^ SC-islets showed no response to the K_ATP_-channel closing sulfonylurea drug glibenclamide (Fig. 1c), providing functional confirmation of channel knockout. As evidenced by increased insulin secretion, glibenclamide effectively closed the K_ATP_-channel of the *KCNJ11*^R201H^ cell lines as expected for the R201H -mutation ^29^. As beta cell hyperactivity in K_ATP_HI is linked to increased oxygen consumption ^32^, we compared the respiration rate of *KCNJ11*^-/-^ and *KCNJ11*^+/+^ SC-islets, finding slightly increased respiration in the former (Fig. 1d). In summary, we were able to recapitulate the expected secretion patterns for all three cell lines, most relevant for the *KCNJ11*^-/-^ SC-islets which showed dramatically increased inappropriate insulin secretion in low glucose. We also identified the optimal timing for maximal phenotypic differences at S7w6, to be used as a test timepoint in subsequent *in vitro* pharmaceutical testing.

### Ghrelin, neuromedin U25 and ropinirole inhibit insulin secretion *in vitro*

After a literature search, we selected seven pharmaceutical agents and hormones that are known to reduce insulin secretion in beta cells but whose effectiveness in the context of K_ATP_HI is unknown. As current K_ATP_HI therapy relies on the SST_2_R-agonist octreotide, we used it as a positive control. For context, we also included exendin 9-39 ^33^, an experimental drug with evidence on effectiveness in K_ATP_HI patients, as well as a GCGR-antagonist. Glucagon analogues are used in the management of CHI due to their insulin counterregulatory, anti-hypoglycaemic effects on the systemic level ^4^. However, on the beta cell level glucagon increases insulin secretion ^34^ prompting our interest in studying the effects of the antagonist. The seven molecules, listed in Table 1, were tested for their ability to modulate the insulin secretion of S7w6 *KCNJ11*^-/-^ SC-islets in 3 mmol/l glucose.

**Table 1.** Pharmaceutical compounds used.

| Name | Mechanism | Supplier | Conc. *in vitro* | Role | Reference |
| --- | --- | --- | --- | --- | --- |
| Acyl-ghrelin, Relamorelin | GHSR agonists | Bachem 4033077<br>MedChem Express HY-19884B | 200 nmol/l<br>500 nmol/l | Candidate molecule | <sup>50</sup> |
| Baclofen | GABA<sub>B</sub>R agonist | Tocris #0796 | 20 µmol/l | Candidate molecule | <sup>57</sup> |
| CYN154806 | SST<sub>2</sub>R inhibitor | Abcam AB141210 | 500 nmol/l | Mechanism testing | <sup>58</sup> |
| DCEBIO | K<sub>Ca3.1</sub> opener | Tocris #1422 | 100 µmol/l | Candidate molecule | <sup>59</sup> |
| Exendin (9-39) | GLP1-R inverse agonist | Gift from Novo Nordisk Indianapolis, Tocris #2081 | 200 nmol/l | Positive control | <sup>33</sup> |
| GIP inhibitor | GIPR antagonist | Gift from Novo Nordisk Indianapolis | 20 nmol/l | Candidate molecule | <sup>60</sup> |
| Glibenclamide | K<sub>ATP</sub>-channel opener | Tocris #0911 | 100 nmol/l | Phenotype validation | <sup>29</sup> |
| L-168,049 | GCGR antagonist | Tocris #2311 | 15 µmol/l | Context | <sup>61</sup> |
| Neuromedin-U25 | NMUR agonist | Bachem 4037521 | 800 nmol/l | Candidate molecule | <sup>62,63</sup> |
| Octreotide | SST<sub>2</sub>R antagonist | Gift from Novo Nordisk Indianapolis | 500 - 800 nmol/l | Positive control | <sup>11</sup> |
| Ropinirole, | D<sub>2-3-4</sub> agonist | Cayman 23871 | 25 µmol/l | Candidate molecule | <sup>64,65</sup> |
| Sumatriptan, | 5-HT<sub>1BD</sub> agonist | Tocris #3586 | 50 µmol/l | Candidate molecule | <sup>66</sup> |

As expected, the positive control agents had a marked inhibitory effect on the insulin secretion of *KCNJ11*^-/-^ SC-islets in 3 mM glucose, approximately 50% for octreotide and 30% for exendin 9-39 (Fig. 2). Of the test agents, GHSR-agonist acyl-ghrelin, NMUR-agonist neuromedin-U25 and D_2_-agonist ropinirole had inhibitory effects in the range of 20-30% (Fig. 2). Based on this *in vitro* testing, we chose the intestinal hormones neuromedin-U25 and acyl-ghrelin for subsequent *in vivo* experiments in mice carrying *KCNJ11*^-/-^ SC-islet grafts.

**Figure 2.**
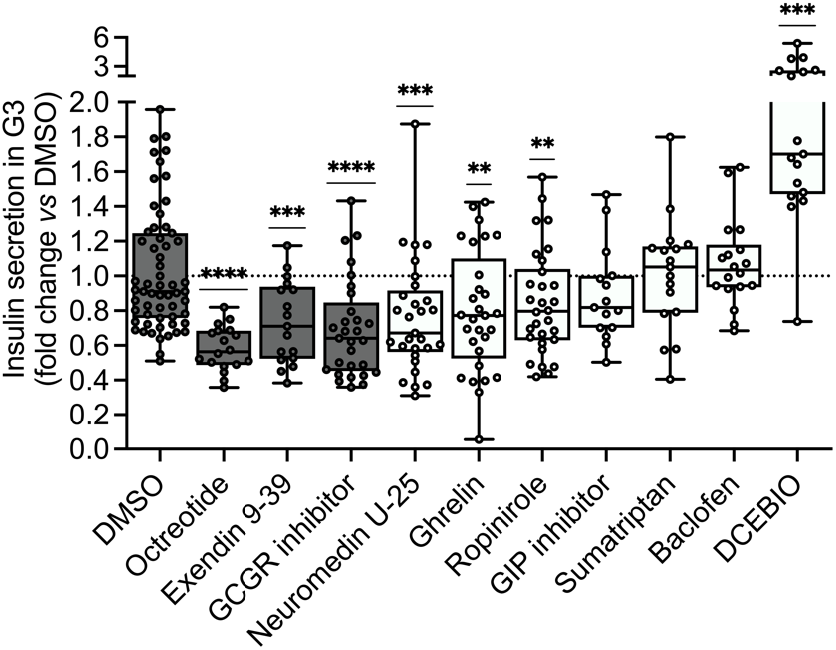
*in vitro* screening for insulin secretion reducing agents: S7w6 *KCNJ11*^-/-^ SC-islets were incubated for 30 minutes in 3 mmol/l glucose in the presence of DMSO or control (dark) or test (white) compounds. Technical replicates shown from N= 6 biologically distinct experiments. Test concentrations and receptors given in table 1.

### Ghrelin receptor agonists increase glycaemia in hypoglycaemic mice implanted with *KCNJ11*^-/-^ SC-islets

We implanted immunocompromised mice with *KCNJ11*^+/+^ or *KCNJ11*^-/-^ SC-islets under the kidney capsule, to create humanised mice where the blood glucose regulation is under the control of the human SC-islets. At 8 weeks after implantation, the *KCNJ11*^-/-^ mice reached human hypoglycaemia in 5-h fasted conditions (average 3.0 mM, 20/27 mice below 3.3mM). The fasting hypoglycaemia persisted until the 16 weeks of the follow-up (Fig. 3a). Meanwhile, the mice implanted with a similar dose of *KCNJ11*^+/+^ SC-islets reduced their blood glucose level from mouse to human euglycaemic level (average 4.2 mM, 4/13 mice below 3.3 mM) (Fig 3a.), highlighting the takeover of blood glucose regulation by the human cells. This rapid and persistent development of hypoglycaemia reproduced the human K_ATP_HI phenotype in mice, more efficiently than in our previous K_ATP_HI SC-islet model^31^ and more severely and persistently than in *Kcnj11*^*-/-*^ mice^22^. Fasting levels of human specific C-peptide, which assess the graft derived insulin secretion, were on average 2-3 times higher in mice carrying *KCNJ11*^-/-^ SC-islet grafts compared to controls. (Fig. 3b).

**Figure 3.**
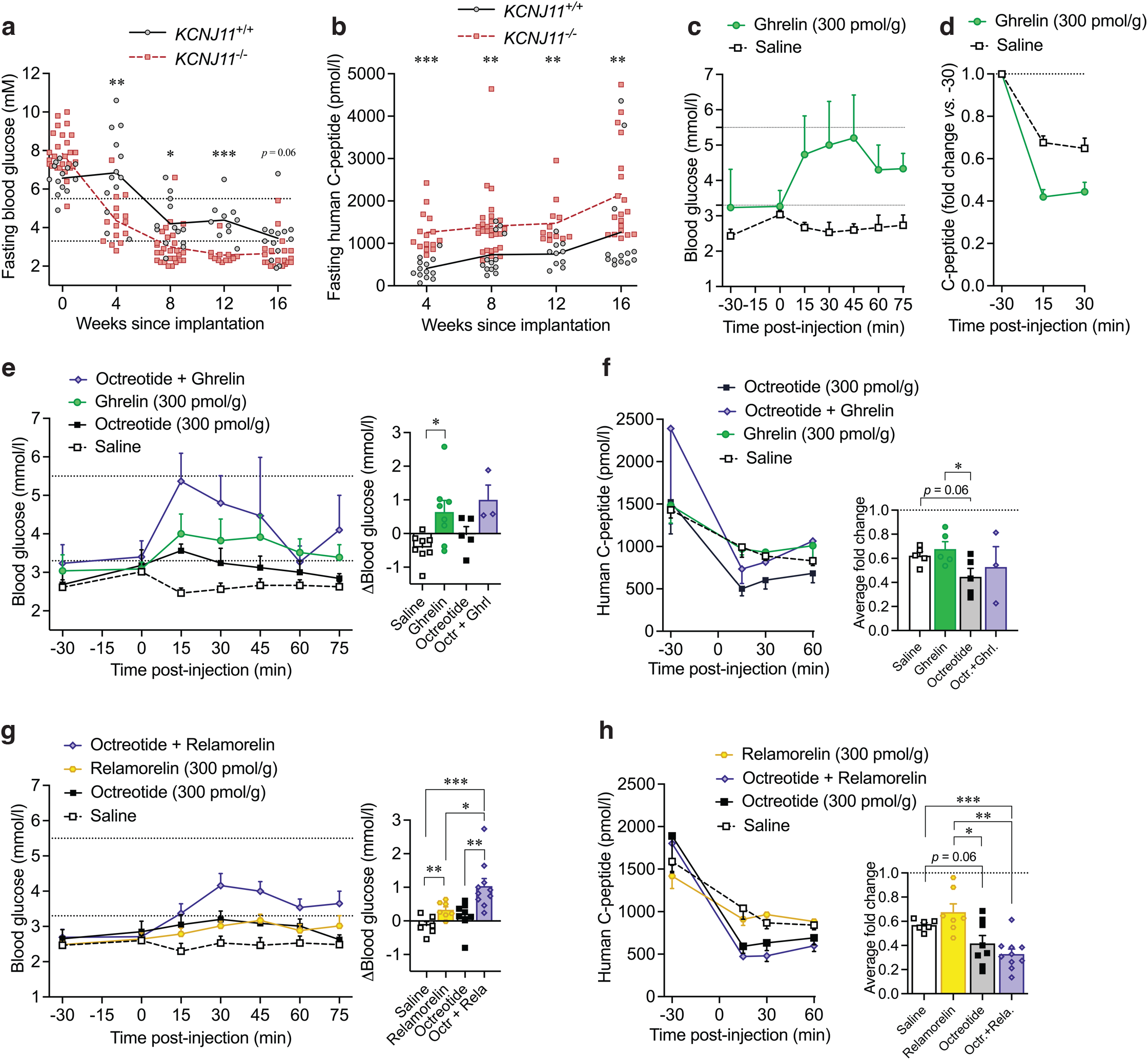
pharmaceutical testing *in vivo*: **a-b)** Glucose level a) and human specific C-peptide level b) after a 5-6 hour fast, in *KCNJ11*^+/+^ and *KCNJ11*^-/-^ SC-islet transplanted mice. Multiple Welch’s t-tests. **c-d)** Glucose level c) and human C-peptide level d) after a 300 pmol/g i.p. injection of acyl-ghrelin at 0 min in *KCNJ11*^-/-^ SC-islet recipients. **e-f)** Glucose level and average change from baseline at 15-75 minutes e) and human C-peptide level and average fold change from baseline at 15-60 minutes f). Data collected after i.p. injections of indicated compounds at 0 min in *KCNJ11*^-/-^ SC-islet recipients. Combination injections consisted of 300 pmol/g of both compounds. **g-h)** Glucose level and average change from baseline at 15-75 minutes g) and human C-peptide level and average fold change from baseline at 15-60 minutes h). Data collected after i.p. injections of indicated compounds at 0 min in *KCNJ11*^-/-^ SC-islet recipients. Combination injections consisted of 300 pmol/g of both compounds. One-way Welch’s ANOVA

To study the candidate molecules *in vivo*, we conducted fasting tolerance tests on mice 8 to 16 weeks after the implantation of *KCNJ11*^-/-^ SC-islets, at the time when they had developed hypoglycaemia. Injections of the pharmaceutical were given to 5-6 h fasted mice 30 minutes after taking basal measurements, to minimize the hyperglycaemic effects of handling stress. Neuromedin-U25 injections increased blood glucose (Fig. S3a) but failed to reduce human C-peptide secretion more than saline. Saline injection inhibited human C-peptide secretion by approximately 30%, presumably through counterregulatory stress hormones released upon experimentation (Fig. S3b). Acyl-ghrelin injections on the other hand both increased blood glucose (Fig. 3c) and reduced C-peptide more than saline injections (Fig. 3d), prompting us to focus on ghrelin for further combination testing.

As octreotide is known to be an effective anti-hypoglycaemic agent for many K_ATP_HI patients, and effective in our SC-islets *in vitro* (Fig 2), we wanted to see how ghrelin compares to its effectiveness and if they have synergistic or overlapping effects. In these combination tests conducted on fasting mice, mice receiving saline had a reduction of blood glucose, which was prevented by octreotide (0.3 mmol/l drop *vs*. 0.0 mmol/l) (Fig. 3e). If ghrelin was administered alone, or in combination with octreotide, blood glucose was increased by appr. 0.5 mmol/l (Fig. 3e). C-peptide secretion was reduced by 40% after saline and ghrelin injections and by 50-60% after octreotide and combination injections (Fig. 3f).

To study this further, we investigated a clinically tested long-acting analogue of ghrelin, relamorelin ^35–38^, in a similar setup in fasting mice engrafted with *KCNJ11*^-/-^ or *KCNJ11*^+/+^ SC-islets. Relamorelin and octreotide both increased glycaemia marginally when administered alone (c. 0.3 mmol/l increase) while mice receiving saline injections exhibited a decrease. However, relamorelin-octreotide co-administration produced a much stronger anti-hypoglycaemic response, more than the sum of each agent alone (1.0 mmol/l increase), reverting the prior hypoglycaemia to human euglycaemia (Fig. 3g). This suggests a synergistic, rather than additive effect. The co-administration reduced human C-peptide secretion in the *KCNJ11*^-/-^ graft-carrying mice by around 70%.

We also studied the effects of relamorelin in a few control mice carrying *KCNJ11*^+/+^ grafts. In them, relamorelin did not change glucose or C-peptide levels more than saline (Fig. S3ef). Saline injections alone were particularly effective in reducing C-peptide in mice carrying *KCNJ11*^+/+^, more so than in mice carrying *KNCJ11*^-/-^ SC-islets (70% *vs*. 40% reduction), suggesting that the latter are less sensitive to physiological regulation by stress hormones (Fig. S3g).

Taken together, ghrelin receptor agonists acyl-ghrelin and relamorelin increase glycaemia and act synergistically with octreotide to restore euglycaemia in hypoglycaemic mice implanted with K_ATP_HI SC-islets.

### Ghrelin receptor agonists reduce insulin secretion of beta cells

While ghrelin’s systemic effects have been well characterised ^39^, they are unknown in the contexts of both K_ATP_HI and stem-cell-derived islets. In our previously collected single cell sequencing data ^27^ integrating primary and SC-islet samples, expression of the ghrelin receptor *GHSR* mRNA is detected mostly in delta cells and to a lesser extent in beta cells (Fig. 4a), but most of the *GHSR* expressing cells come from the integrated primary islet samples. This expression profile, consistent with other reports ^40^, could indicate that ghrelin receptor agonists act primarily through the delta cells by increasing somatostatin secretion which would in turn decrease insulin secretion, shown to be the major mechanism in mice ^41,42^.

**Figure 4.**
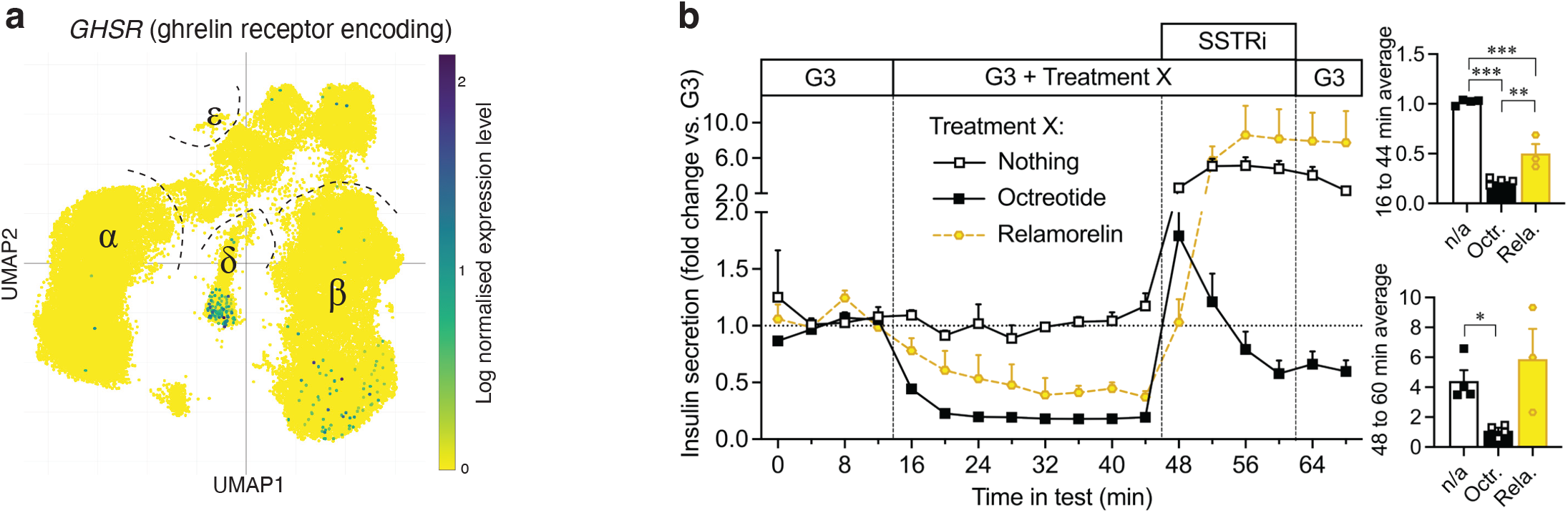
GHSR *in vitro* expression and activity: **a)** Single cell mRNA sequencing profile for the expression of the ghrelin receptor gene *GHSR* in *KCNJ11*^+/+^ primary and SC-islet endocrine cells, cell types indicated with Greek lettering, from Balboa, Barsby & Lithovius et al. data, *Nat. Biotech 2022*. **b)** Dynamic perifusion on S7w9 *KCNJ11*^-/-^ SC-islets, treated with 3 mmol/l glucose (G3), 500 nmol/l octreotide, 500 nmol/l relamorelin and 500 nmol/l CYN154806, a somatostatin receptor antagonist (SSTRi). Insets display fold change averages at two phases of the test. One-way Welch’s ANOVA.

To study this experimentally, we challenged SC-islets with octreotide (SSTR agonist) and relamorelin with and without co-administration of a SSTR antagonist CYN154806, to detect potential competition for the same receptor (Fig. 4b). Octreotide and relamorelin both inhibited insulin secretion in *KCNJ11*^-/-^ islets, verifying again that both are active on the islet level. To our surprise, the SSTR antagonist produced very strong secretory stimulation in otherwise untreated SC-islets, indicating high levels of paracrine inhibition of insulin secretion by the delta cells in the *KCNJ11*^-/-^ SC-islets (Fig. 4b). When added after octreotide, the stimulation by SSTR antagonist was less strong, indicating the expected competition for the same SSTR by both agents. When SSTR antagonist was added after relamorelin, relamorelin was not able to reduce the stimulatory effects of SSTR antagonist (Fig. 4b), which can be due to its lower overall ability to suppress beta cell insulin secretion. These results indicate that the beneficial blood glucose increase of relamorelin we detected *in vivo* is not mediated only by acting on the systemic level, but also on the islet level, where the major mechanism likely is to increase somatostatin secretion from delta cells.

## DISCUSSION

In this study, we used an optimal model of K_ATP_-channel related congenital hyperinsulinism, human *KCNJ11*^-/-^ SC-islets implanted in mice, to identify glycaemia restoring effects of ghrelin-receptor agonists. We demonstrate that long-acting ghrelin-receptor agonist has a synergistic effect with octreotide, resulting in reversal of hypoglycemia.

More specifically, we show that administration of ghrelin or relamorelin induces an increase in blood glucose in 15 minutes, which is sustained for at least 75 minutes, despite continued fasting of the mice. This setup rules out the possibility that these drugs would act by inducing food-seeking behaviour, rather pointing to effects on the islets and insulin sensitive organs. The relamorelin-octreotide combination therapy demonstrated more potent effects on blood glucose than octreotide alone, although both treatments had similar effects on C-peptide levels. This suggests that relamorelin’s added benefit on glycaemia is primarily mediated through systemic effects on insulin-sensitive organs. While relamorelin does act on the islets, it is likely not the primary mechanism of action in this combination therapy *in vivo*. Given this mechanism of action, relamorelin-octreotide therapy would plausibly also be effective in most of the other, rarer forms of CHI besides K_ATP_HI. Particularly those forms where increased activity of the triggering pathway of insulin secretion is the central concern, e.g. forms caused by *HK1* and *GCK* mutations ^43–45^ are mechanistically similar to K_ATP_HI tested here.

Ghrelin, a hormone secreted by P/D1 enteroendocrine cells in the stomach ^46^, as well as pancreatic epsilon cells ^47^, has been linked to regulation of many physiological processes, such as the feeling of hunger, secretion of adrenocorticotropic and growth hormones, augmentation of cardiac output and enhancement of intestinal mobility ^39^. We included it for testing for its known role in increasing blood glucose and decreasing insulin secretion in human *in vivo* ^48,49^ and suppressing insulin secretion in human islets *in vitro* ^50^ in the context of healthy islets. Ghrelin’s systemic glucoregulatory effects are mediated by increases in lipolysis and increases in cortisol and growth hormone secretion, both of which have effects opposing those of insulin ^39,49^. In beta cells, the activation of GHSR leads to activation of inhibitory G-proteins (G_αi_) that increase repolarising currents from the Kv-channels and ultimately attenuate Ca^2+^ dependent insulin secretion ^51^, at least in rat. These effects reduce insulin secretion by suppressing both the triggering and amplifying signals for insulin release. Additionally, in mice, ghrelin is a potent stimulant of delta cell somatostatin secretion resulting in SSTR-mediated suppression of insulin secretion on the beta cell ^41,42^. Similar mechanisms are likely active in humans, as GHSR expression is predominantly observed in delta cells ^40^.

Natural acyl-ghrelin has a half-life of <10 minutes, which has prompted the development of long-acting recombinant analogues such as relamorelin (half-life ranging from 4.5 to 19 hours) ^37,38^. Relamorelin, also known as RM-131, has been clinically tested for anorexia nervosa ^52^ (NCT01642550), and more recently, in >650 type 1 and 2 diabetes patients experiencing diabetic gastroparesis (NCT01571297, NCT03383146 ^35,36^. Relamorelin was found to increase gastric motility ^36^ and received FDA fast track designation for gastroparesis. However, its development was terminated before it received market authorisation due to economic reasons. The main side effects in the gastroparesis studies were related to increases in blood glucose and worsening of type 2 diabetes control, increasing fasting glucose by 3 mmol/l in the group receiving the highest dose ^35^. This is likely mediated by detected decreases in C-peptide and increases in cortisol and growth hormone levels ^38^. Otherwise, relamorelin has been well tolerated. This side effect profile makes relamorelin an attractive candidate drug for K_ATP_HI. Speculatively, in a combination therapy with octreotide for K_ATP_HI, the effects of relamorelin in increasing growth hormone levels and gastric motility could counteract the growth hormone reducing ^53^ and gastric motility decreasing ^54^ effects of octreotide. These are clinically relevant side effects in CHI, as high dose octreotide therapy has been linked to growth restriction ^53^ and necrotising enterocolitis ^16,17^, that is partially mediated by decreased intestinal motility ^55^.

We found that SSTR antagonism greatly stimulates insulin secretion in low glucose conditions both in *KCNJ11*^+/+^ and *KCNJ11*^-/-^ SC-islets. This would indicate that under sub-stimulatory conditions, paracrine somatostatin signalling has a substantial negative effect on insulin secretion in human SC-islets. This basal somatostatin signalling was recently reported in mice by Huang and colleagues ^56^. Nevertheless, even with the basal somatostatin signalling, marked further suppression can clearly be achieved with octreotide. We also demonstrate that on the beta cell level, GCGR antagonism results in decreased insulin secretion in *KCNJ11*^-/-^ SC-islets, which would be beneficial in the K_ATP_HI context, while on the systemic level GCGR agonists are used.

As mentioned, the GHSR agonists act both on the islet and on the systemic level. In our system, the *KCNJ11*^-/-^ islets are human and the GHSR agonists’ effects on the islets can be expected to translate directly to K_ATP_HI patients. However, this is not necessarily the case for the systemic effects of GHSR agonists which are mediated by murine organs in our system. This represents the major weakness in our study. As described, GHSR agonists are known to increase blood glucose in the human context, mitigating this concern.

In conclusion, we have identified ghrelin receptor agonists as insulin secretion reducing agents that induce normoglycaemia-restoring effects in a stem cell derived islet-based model of K_ATP_HI. As such, we propose testing relamorelin-octreotide combination therapy in K_ATP_HI patients to improve pharmaceutical management of this disorder that remains challenging to manage with the currently available means.

## METHODS

### Differentiation of stem-cell-derived islets

Human embryonic stem cell line H1 genome-engineered for homozygous knockout of the *KCNJ11* gene, were used as the K_ATP_HI cells, as we have previously described ^28^. Unedited H1 cells were used as isogenic controls. Stem cell maintenance and differentiation experiments were carried out as described in our 2022 protocol ^30^, except that the SC-islets were transferred to suspension after 48h in the microwells.

### *In vitro* insulin secretion tests

For phenotypic validation (Fig. 1), sequential static incubations were performed as we have previously described ^27^. For the small-scale screening (Fig. 2), 1000-2000 S7w5-w7 SC-islets were picked on a separate plate and washed twice with Krebs-Ringer buffer with 3 mmol/l glucose (KRB). 15-20 SC-islets were then picked per each well of a 96-well plate in 100 μl KRB, with 11-12 replicates for the controls, DMSO treated *KCNJ11*^-/-^ SC-islets and 6 replicates for the experimental conditions. 150 μl KRB was added on top of each well and incubated in rotation until 2h had elapsed since the first washes with KRB. The SC-islets were then washed once by removing, adding, and removing again 200 μl of KRB. 200 μl of KRB containing the test compounds was then added and SC-islets incubated for 30 min in rotation, after which 50 μl sample was taken on another 96-well plate and both plates frozen. The time elapsed during addition of the final wash medium, the test compounds as well as during pipetting the secretion samples were recorded, and the final secretion values were corrected for differences caused by the different duration of these steps for individual wells and then normalised for DNA content of each well.

Dynamic perifusion experiments (Fig. 4) were conducted on PERI-4.1 apparatus (Biorep Diabetes) on 50 S7w9 SC-islets that were picked for each test on the previous day to 12-well plates in S7 culture medium. 1.5-hour acclimatation at 3 mmol/l glucose preceded the test. All insulin secretion samples were analysed on insulin ELISA kits (Mercodia) per manufacturer instructions. Test compound used in the tests, as well as their suppliers and concentrations are listed in **table 1**.

### Respirometry

Basal respiration rate was determined in 3 mmol/l glucose in stage 7 culture medium from 8 technical replicates, using the SeaHorse XF96 analyser, as previously described ^27^.

### *In vivo* experiments

Male NSG mice were implanted under the kidney capsule with 3-6 mm^3^ of *KCNJ11*^+/+^ or 4-5.5 mm^3^ *KCNJ11*^-/-^ SC-islets (1 mm^3^ = 566 islet equivalents c. 150 SC-islets), as previously described ^27^, with preimplantation dose quantification using CellProfiler pipeline to individually quantify the volume of the SC-islets in each preparate to be implanted as previously described ^28^. They were housed in 12 h/12 h light-dark cycle and fed irradiated standard chow. All follow-up blood samplings as well as pharmaceutical tests were conducted on 5-6 h fasted animals in the early afternoon at 8 to 16 weeks post-implantation. Mice were weighted and sampled for basal measurements 30 minutes before test molecule injections, which were given in 10 μl/g of 0.9% NaCl intraperitoneally. Blood glucose and samples were taken at -30, 15, 30 and 60 min, and simple blood glucose measurements on 0, 45 and 75 min. Each mouse was subjected to a different test condition at each test, with maximum 3 tests performed on each mouse.

### Statistical methods

Analyses were performed in GraphPad Prism v9. Data points are biological replicates (individual SC-islet batches or individual animals) and composed of multiple technical replicates unless otherwise indicated. *P*-values are shorthanded as <0.05 *, <0.01 **, <0.001 *** and statistical tests used are indicated in figure legends.

## ACKNOWLEDGEMENTS

Jarkko Ustinov and Solja Eurola (University of Helsinki) are thanked for technical assistance. Dr. Vikash Chandra and Eliisa Vähäkangas (University of Helsinki) are thanked for critical comments. Dr. Brian Finan (Novo Nordisk Indianapolis research site) is thanked for supplying octreotide, exendin 9-39 and the GIP-antagonist. This research was funded by the Academy of Finland (Center of Excellence “Metastem”, grant #312437), Sigrid Jusélius foundation and Diabetestutkimussäätiö. VL gratefully acknowledges personal grant support from Emil Aaltosen säätiö, Ida Montinin säätiö, Diabetestutkimussäätiö, Maud Kuistila memorial foundation, Suomen Lääketieteen säätiö, Biomedicum Helsinki foundation and Orion research foundation for this project and others.

## Author contributions

VL designed the study and chose the candidate molecules, conducted SC-islet differentiations and *in vitro* tests, assisted in animal experimentation, analysed and visualised data and wrote the manuscript. HM and JSV conducted animal experimentation and edited the manuscript. HI generated the *KCNJ11* cell lines and edited the manuscript. TB conducted respirometry and edited the manuscript. DB processed transcriptomics data and edited the manuscript. TO conceptualised and supervised the study, edited the manuscript and provided resources. All authors agree on the publication of this work.

