## Supplementary material for "Ghrelin and relamorelin alleviate hypoglycaemia in humanised mice with congenital hyperinsulinism": Figure 3 supplement

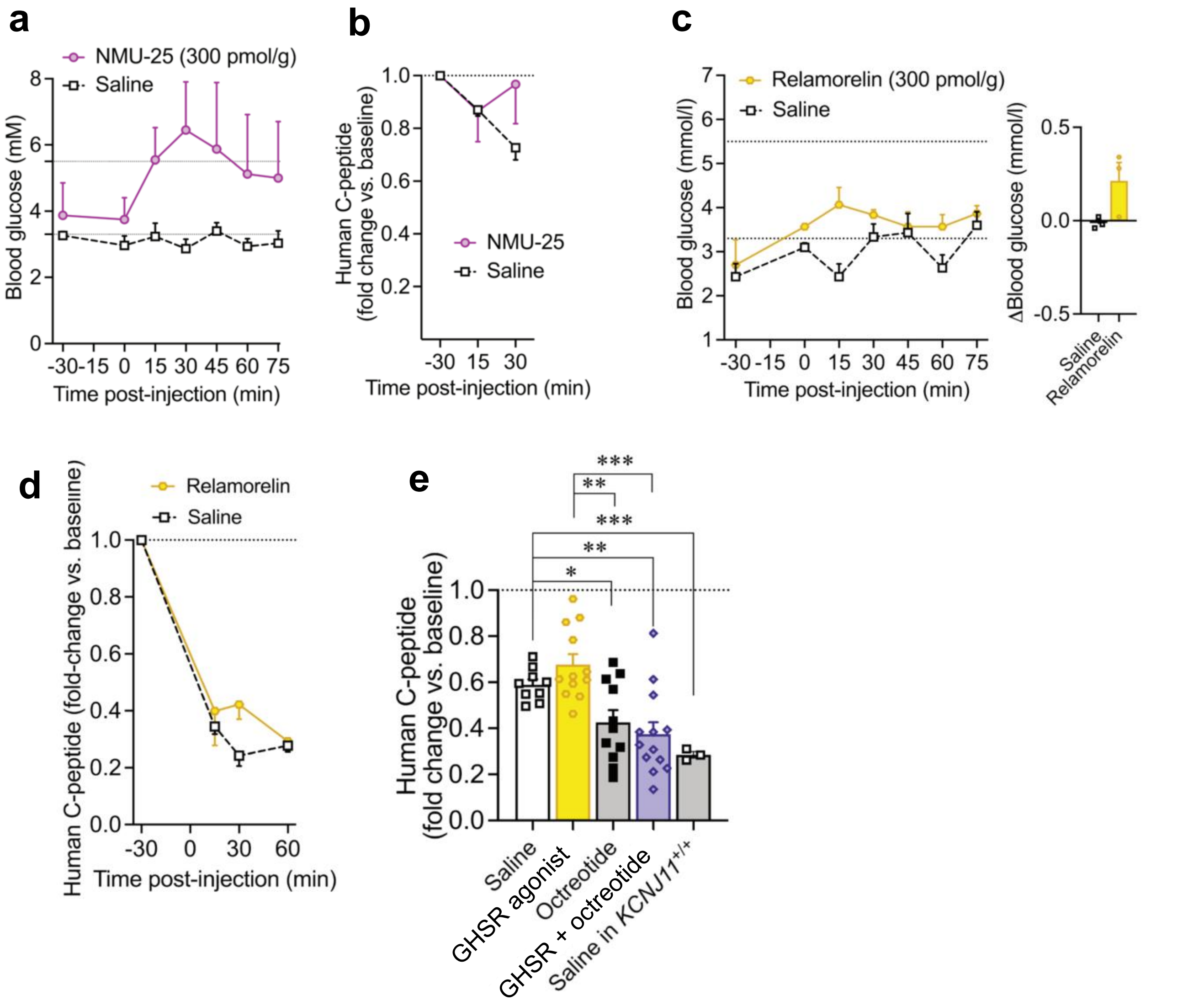


**Figure 3 supplement: a-b)** Glucose level a) and human C-peptide level b) after a 300 pmol/g i.p. injection of neuromedin-U25 at 0 min in *KCNJ11*$^{-/-}$ SC-islet recipient cohort 1. **c-d)** Glucose level and average change from baseline at 15-75 minutes c) and human C-peptide fold change from baseline d) in *KCNJ11*$^{+/+}$ recipients. Data collected after i.p. injections of indicated compounds at 0 min **e)** Average C-peptide fold change from baseline at 15-60 minutes. Pooled data from tests using acyl-ghrelin and relamorelin (GHSR agonists) on *KCNJ11*$^{-/-}$ recipients shown separately in Fig. 3f and 3h, with *KCNJ11*$^{+/+}$ recipients added for context. One-way Welch's ANOVA.
